# A leaf-expressed *TERMINAL FLOWER1* ortholog from coffee with alternate splice forms alters flowering time and inflorescence branching in *Arabidopsis*

**DOI:** 10.1101/2024.08.14.607758

**Authors:** Carlos Henrique Cardon, Victoria Lesy, Catherine Fust, Thales Henrique Cherubino Ribeiro, Owen Hebb, Raphael Ricon de Oliveira, Mark Minow, Antonio Chalfun Junior, Joseph Colasanti

## Abstract

Perennial, polycarpic species, such as *Coffea* sp L. (coffee), exhibit asynchronous flowering while maintaining concomitant vegetative growth. This growth dichotomy is associated directly with fruit development and maturation time. To identify molecular components that underlie asynchronous flowering, we isolated phosphatidylethanolamine binding protein (PEBP) homologs expressed in coffee and identified a gene with high similarity to Arabidopsis *TERMINAL FLOWER1-like*. In Arabidopsis, interaction of TFL1 (AtTFL1) with bZIP transcription factor floral regulator FD (AtFD) forms a floral repressor complex that maintains inflorescence meristems in an indeterminate state. Unlike *AtTFL1*, which is expressed only in the shoot apical meristem, *CaTFL1* transcript was detected exclusively in coffee leaves. Moreover, this transcript retained an intron, which was not reported for *AtTFL1*. CaTFL1 was characterized through heterologous expression in *Arabidopsis* and protein interaction analysis. Ectopic overexpression of *CaTFL1* in transgenic *Arabidopsis* plants caused extreme late flowering or prevented flowering. However, the most severe floral repressive activity occurred in transgenic plants that spliced out the extra intron from *CaTFL1*. Yeast two hybrid assay revealed that CaTFL1 protein encoded by the spliced mRNA interacts with AtFD, as well as *Arabidopsis* 14-3-3 protein. These findings suggest that *CaTFL1* acts as a leaf-expressed floral repressor, whose activity is controlled by alternate splicing, and may contribute to asynchronous flowering in coffee.

## INTRODUCTION

Plants endure constant challenges to maintain optimal vegetative and reproductive growth stages due to seasonal variations and changes in climate. Whereas successful reproduction is crucial for the propagation and evolution of plant species, sufficient vegetative biomass is also important for plant survival. Many regulatory networks help plants respond to varying conditions such as temperature, water stress, disease, insect damage, photoperiod, and other environmental factors in order to balance these two phases and optimize growth.

In annual angiosperms, such as *Arabidopsis thaliana*, the transition from vegetative to reproductive growth is strongly affected by day length (photoperiod) (Albani et al., 2010; Amasino, 2010). The transition from vegetative to reproductive growth occurs when the shoot apical meristem (SAM) produces inflorescence meristems (IM) from which determinate floral meristems (FM) form, giving rise to flowers. Maintenance of indeterminate IMs and the production of determinate FMs specify the reproductive architecture of flowering plants, including lateral inflorescence branches. The balance of inflorescence branch formation and the amount of FM producing terminal flowers is regulated by the interplay of members of the phosphatidylethanolamine binding protein (PEBP) super family with opposing activities. In *Arabidopsis* these are represented by *TFL1* and *FLOWERING LOCUS T* (*FT*), the latter of which encodes a florigen (Ahn et al., 2006; Conti & Bradley, 2007; Corbesier et al., 2007). Together these *PEBP* genes encode related proteins that compete to regulate the balance of indeterminate IMs to determinate FMs. Despite their antagonistic functions, these proteins are highly conserved in flowering plants, sharing highly conserved amino acid sequence motifs (Moraes et al., 2019; Wang et al., 2018).

The floral induction and shoot architecture roles of highly conserved PEBP superfamily *FT/TFL1* genes from different plant species have been studied through heterologous expression in *Arabidopsis* (Coelho, Minow, Chalfun-Júnior, & Colasanti, 2014; Lazakis, Coneva, & Colasanti, 2011). *TFL1-like* genes include the floral repressors *TERMINAL FLOWER 1* (*TFL1*), *BROTHER* of *FT* (*BFT*) and *CENTRORADIALIS* (*CEN*), whereas floral inducers include *FLOWERING LOCUS T* (*FT*), *MOTHER OF FT AND TFL1 (MFT)* and *TWIN SISTER* of *FT* (*TSF*) (Goretti et al., 2020; Yamaguchi et al., 2005; Yoo et al., 2010). *Arabidopsis TFL1* is expressed in inflorescence meristems where it is required to maintain an indeterminate state. Competition between TFL1 and FT for interaction with bZIP transcription factor, FD, specifies the level of indeterminacy (Hanano & Goto, 2011; Jaeger, Pullen, Lamzin, Morris, & Wigge, 2013). The floral repressor complex (FRC) consists of a complex of TFL1 and FD plus 14-3-3 proteins, whereas the floral activation complex (FAC) forms when FT interacts with FD and 14-3-3-proteins. The FAC irreversibly promotes the expression of flower identity genes such as *APETALA 1* (*AP1*), *LEAFY* (*LFY*), and *FRUITFULL* (*FUL*) to form determinate flowers (Conti & Bradley, 2007; Kaneko-Suzuki et al., 2018).

In annual flowering species, differentiation of vegetative meristems to reproductive meristems in the same year herald a shift to the end of their life cycle. Conversely, polycarpic perennial species must maintain vegetative growth concomitantly with flower induction year after year, suggesting more complex control of meristem identity (Foster et al., 2003). The role of *TFL1* in annual species is well documented in *Arabidopsis* (Moraes et al., 2019), however, much less is known about *TFL1* orthologs in perennial species (Bustin et al., 2009; Klocko et al., 2016; Mimida et al., 2009; Mohamed et al., 2010). In perennial species, the underlying mechanisms that coordinate formation of determinate and indeterminate SAMs, axillary meristems (AM) and the transition from vegetative to reproductive development require further study.

A better understanding of the reproductive/vegetative balance in perennial plants may shed light on how asynchronous flowering is regulated. Asynchronous flowering in perennials, such as *Coffea* sp L., results in uneven fruit maturation, which affects fruit harvest and, ultimately, beverage quality. *C. arabica* is an allopolyploid that originated from interspecific crossing between *C. canephora* and *C. eugenioides* (Carvalho, 1946). Coffee plants form a central stalk with lateral growing (plagiotropic) and upward growing (orthotropic) branches that give rise to FMs derived via axillary meristem (AM) differentiation. The dispersal of floral buds results in the distribution of coffee fruit all along the branch (Davis Fls, Govaerts, Fls, & Stoffelen, 2006).

Given the conserved role of *TFL1*-related genes in controlling shoot architecture and branch patterning in diverse plant species, we investigated a putative coffee *TFL1* ortholog through *in silico* analysis, heterologous expression in *Arabidopsis* and protein interactions. Our findings suggest that *CaTFL1* is a component of a floral repressor complex (FRC) that regulates inflorescence architecture and flowering time. Unexpectedly, we also discovered that *CaTFL1* has spliced and unspliced leaf transcript variants. Whereas the spliced *CaTFL1* mRNA encodes an active repressor protein, the unspliced version lacks repressive activity, suggesting a post-transcriptional control mechanism for this regulatory gene that specifies coffee floral induction and shoot architecture.

## RESULTS

### Identification of a *TFL1* homolog in *Coffea sp*

Isolation and local alignment of PEBP gene sequences from public coffee sequence databases revealed several homologs with close homology to *TFL1-like* genes from other plant species (**Fig. 1A, Fig. S1**). Multiple global alignments and phylogenetic analyses of these sequences revealed putative *C. arabica TFL1* genes based on similarity with known protein sequences in *Arabidopsis thaliana* and *Solanum lycopersicum* that have well-established roles in regulating flowering and shoot architecture (Prusinkiewicz, Erasmus, Lane, Harder, & Coen, 2007). We identified three putative *TFL1*-related genes in *C. arabica,* including one from each parental line and putatively duplicated loci on chromosomes 1 (ch1), ch2 and ch6. Since these sequences have (**Fig. S1**) amino acid identities above 93%, we selected the *C. arabica* ch2 sequence, designated *CaTFL1*, for further analysis. The similarity of this putative coffee *TFL1* ortholog to other *TFL1*-like PEBP genes (**Fig. 1A**) suggests that it might be involved in flowering repression rather than floral induction (Jin et al., 2020). Also, *CaTFL1* contains conserved amino acids, such as residues R63, P95, F102 and R132 and H/Y85, which have been shown previously to mediate FRC/FAC docking via 14-3-3 protein (**Fig. 1B**).

**Figure 1.**
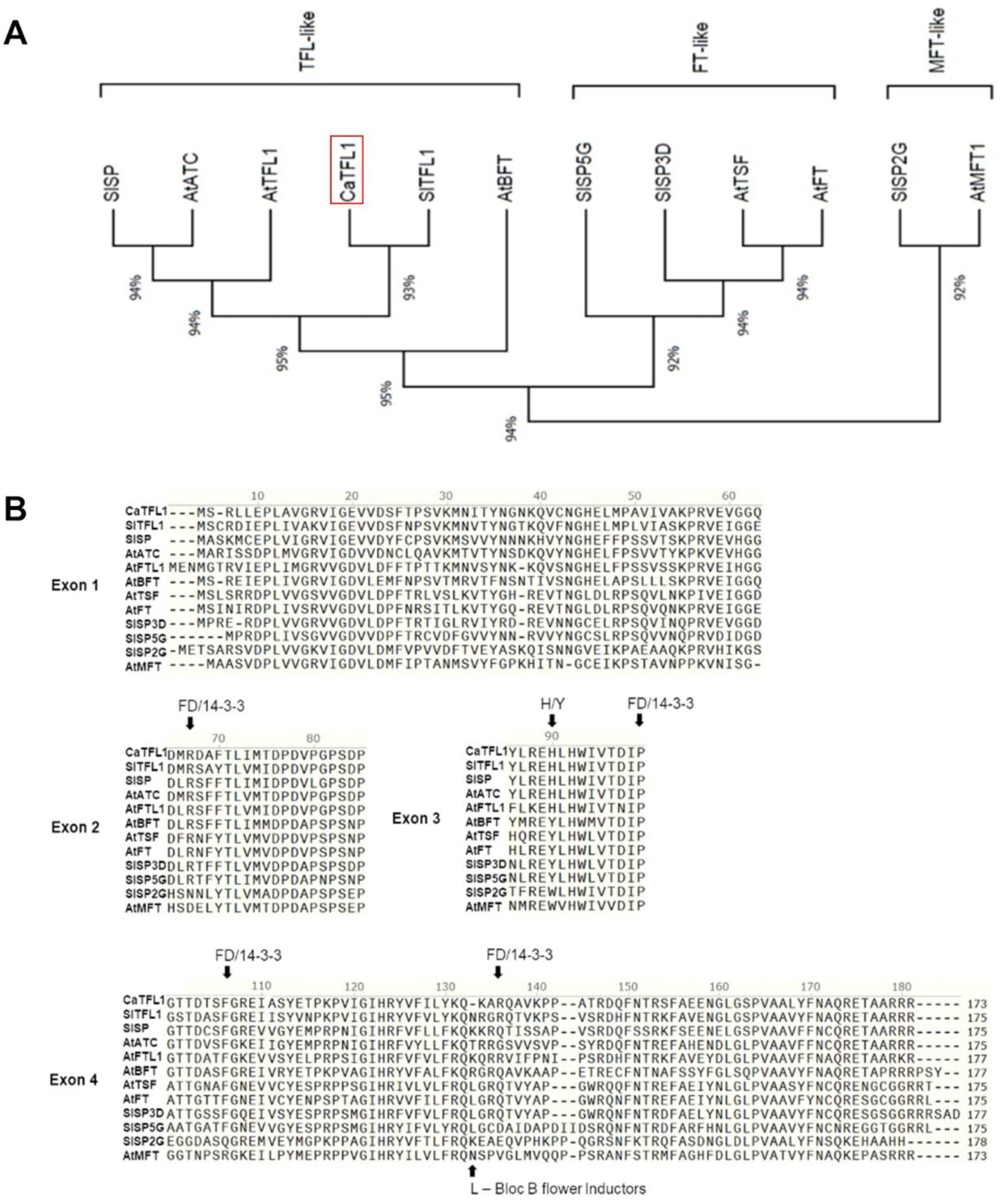
CaTFL1 amino acid sequence grouped with PEBPs members from *Arabidopsis thaliana (At)* and *Solanum lycopersicum (Sl*). **A** – CaTFL1 (Highlighted in red) grouped in the TFL-like group through phylogenetic tree formed by the three main PEBP flower control genes, TFL-like, FT-like and MFT-like. **B** – *At* and *Sl* PEBPs amino acid sequences aligned with CaTFL1 amino acid sequence showing residues potentially involved in FD and 14-3-3 complex formation in each exon sequence and, responsible for separate the TFL-like group from FT-like and MFT-like group. CaTFL1 is placed in the first alignment line.

### *CaTFL1* transcripts isolated from coffee leaves retain an intron

In *Arabidopsis, TFL1* mRNA is expressed and translated in the SAM early in development, where it competes with FT for interaction with the FD/14-3-3 complex (Conti and Bradley, 2007; Goretti et al., 2020). However, the *CaTFL1* transcript examined here was isolated and amplified from coffee leaf, where it is predominantly expressed. The gene structures of *Arabidopsis TFL1* and *CaTFL1* are similar, with each having four exons and three introns (Fig. S2). Unexpectedly, all mature mRNA sequence isolated as cDNAs from coffee leaf retained the third intron, whereas the other three introns were spliced out, similar to orthologous *TFL1* genes from *Arabidopsis* and other species (Fig. S2). The amplified fragment was 689 bp, of which 522 bp corresponded to the deduced *CaTFL1* sequence. The remaining 167 bp corresponded to intron 3 retained in the mature mRNA. Only the 689 bp fragment generated by PCR from coffee leaf cDNA was detected (**Fig. S2**), suggesting that the unspliced version is the dominant transcript in leaf tissue. The transcript that retains intron 3 is designated *as-CaTFL1*, whereas the completely spliced version that encodes a protein highly similar to TFL1 proteins from other species is designated *s-CaTFL1*.

Since these samples were taken at the same time of year in autumn (April), we examined whether different splice variants were found at different times of the year. Transcripts from two *C. arabica* genotypes, Iapar 59 and Acaua, were amplified from leaf samples taken in December, February, April, June and October. December in Brazil has the longest days (13.5 hours) while June has the shortest days (9 hours). RT-PCR showed that the larger 689 bp cDNA was the only product detected in most months in both of these genotypes (Fig. 2). However, the completely spliced version (522 bp) of *CaTFL1*, corresponding to *s-CaTFL1,* was the main transcript detected in leaf tissue taken in June, or winter in Brazil (**Fig. 2**). Moreover, transcript levels from June samples were reduced relative to *as-CaTFL1* transcripts from other times of the year. It should be noted that in the coffee genotype, Conilon, only as-*CaTFL1* transcripts were detected, and only in June and October (Fig. 2), implying a different mode of CaTLF1 splicing in the Conilon branches sampled.

**Figure 2.**
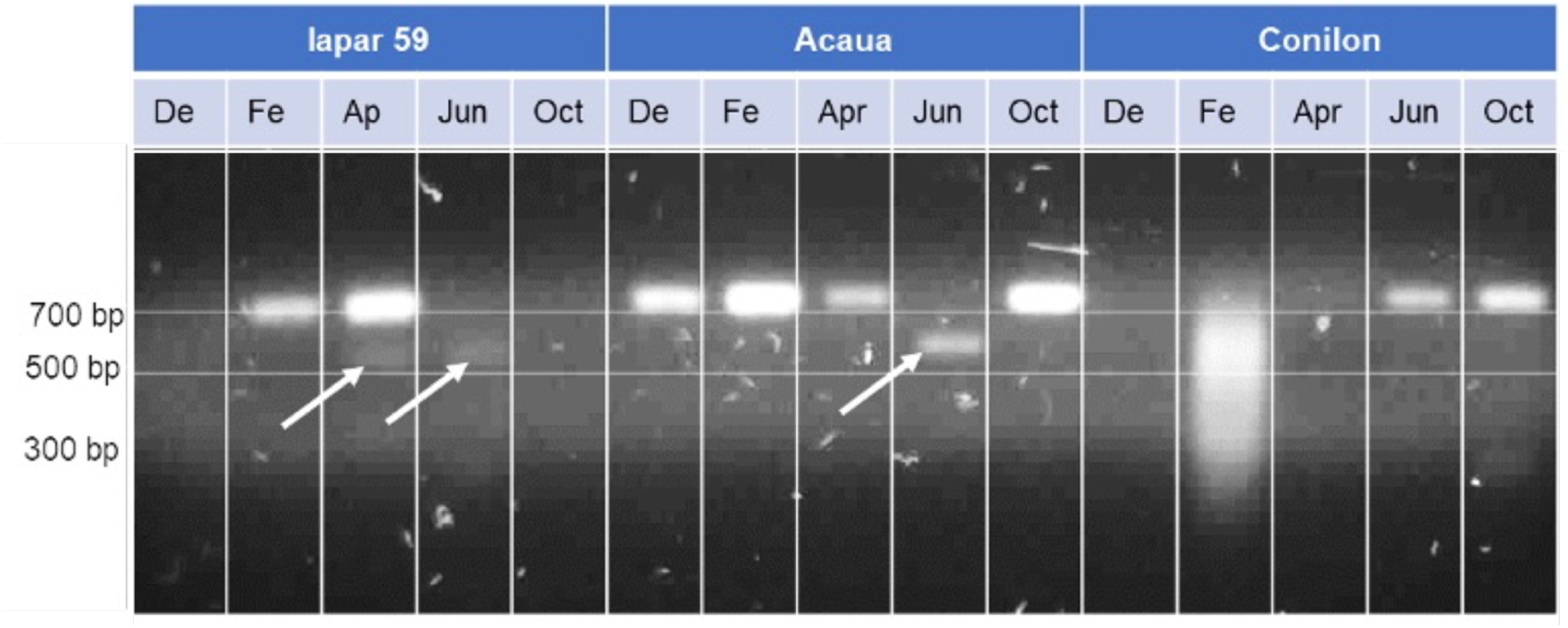
*CaTFL1* transcript amplified from two *Coffea arabica* genotype (Iapar 59 and Acauã) and, one *Coffea canephora* genotype (Conilon) in five different time of the year (De – December, Fe – February, Ap – April, Jun – June and, Oct – October). On the left side are identified the ladder length compared with the PCR amplicons.

To further examine the *CaTFL1* leaf transcript pattern we performed an alignment with RNAseq libraries from coffee leaves available in the NCBI database comparing the *C. arabica* and *C. canephora* genomes (**Fig. 3**). Interestingly, the aligned reads against the coffee genome showed that the unspliced leaf version was only in the *C. eugenioides* subgenome *CaTLF1* copy within the *C. arabica* genome (**Fig. 3A**), with no reads aligning to the retained intron over the *C. canephora* sub genome (**Fig. 3B**). The same was observed for reads from *C. canephora* leaf *CaTFL1* transcripts aligned against the *C. canephora* genome, in which no intron regions were retained in the mature transcripts (**Fig 3C**).

**Figure 3.**
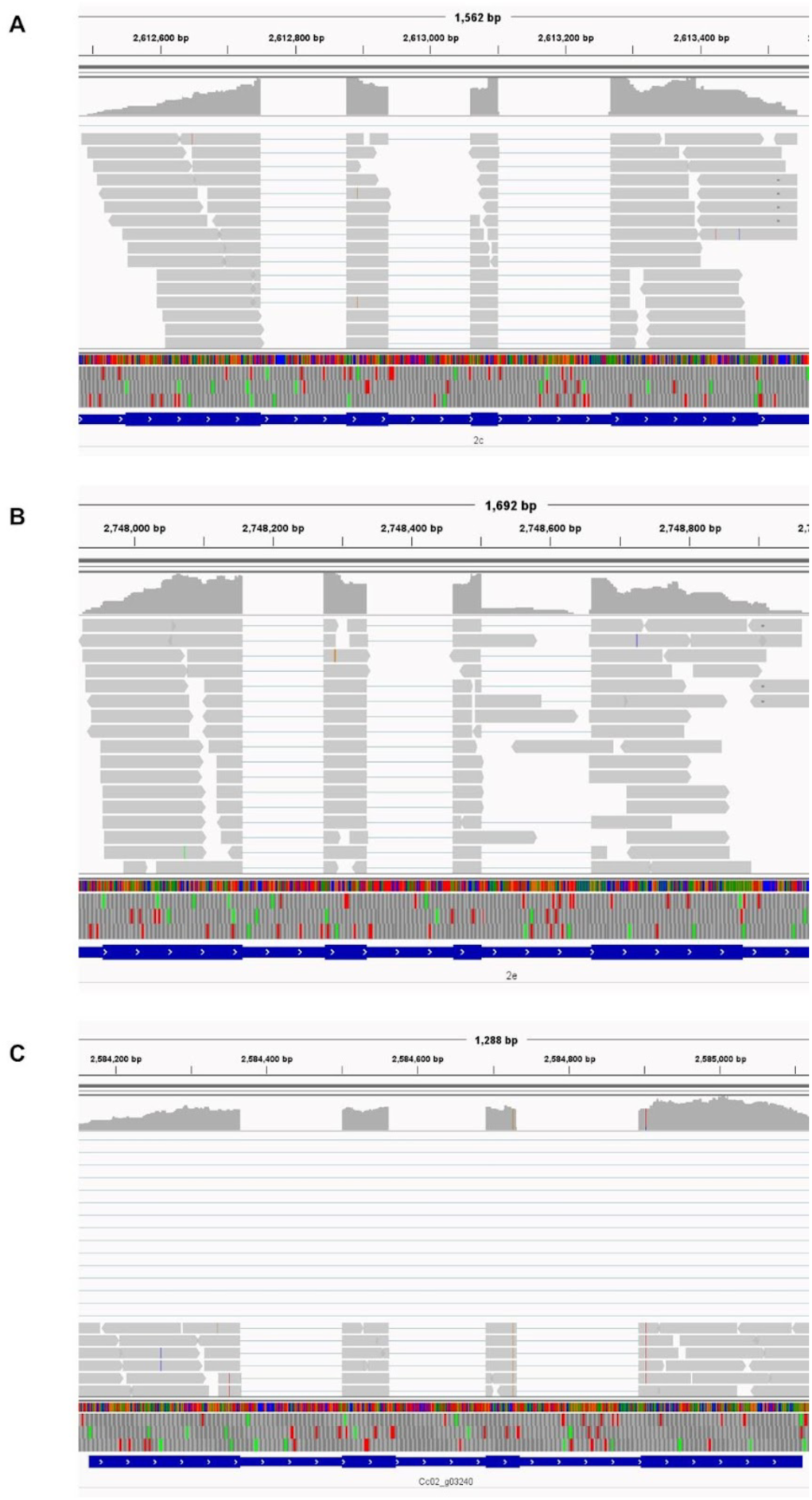
Interactive Genome View (IGV) analysis of an intron retained in mature *CaTFL1* mRNA in the *Coffea eugenioides Coffea arabica* genome subversion in transcript RNAseq alignment from *C. arabica* leafy sample libraries. Gray bars in the figure indicate de reads aligned with the genome showed in the inferior part, followed by a blue bar that represent the CaTFL1 mRNA sequence divided in four exon blocs A - *CaTFL1* alignment in *Coffea canephora* chromosome 2 subversion sequence in *Coffea arabica* genome. **B** - *CaTFL1* alignment in *Coffea eugenioides* chromosome 2 subversion sequence in *Coffea arabica* genome. **C** – *CcTFL1* alignment in *Coffea canephora* genome with transcripts from *Coffea canephora* leafy sample libraries.

### Overexpression of as-*CaTFL1* in *Arabidopsis* causes late flowering and excessive branching

The *as-CaTFL1* cDNA isolated from coffee, which retained intron 3, was over-expressed in *Arabidopsis* under the control of the *CaMV 35S* promoter to test its effect on flowering and plant architecture. All first generation (T1) *CaTFL1* overexpression transgenic *Arabidopsis* lines displayed strong floral repression, with increased rosette leaf number, and weak apical dominance that altered the IM branching architecture in both WT and *tfl1-14* mutant backgrounds (**Fig. 4**). Transgenics in the latter background show that ectopic *CaTFL1* expression could restore indeterminate growth in a *tfl1* mutant. Indeed, ectopic *CaTFL1* expression caused inflorescence aberrations by altering central inflorescence branch development and increasing inflorescence branch numbers. With regard to flowering time and rosette leaf numbers, overexpression of *CaTFL1* was more severe in WT Col-0 than in the *tfl1* mutant background. Comparison of 9 independent T1 lines showed that all flowered later than 9 *tfl1* T1 lines with the *35S::CaTFL1* transgene. Representative transgenic plants in the Wt and mutant background are shown in **Fig. 4A** and **4B**, respectively. Overexpression of *AtTFL1* in WT *Arabidopsis* (Col-0) also caused late flowering and increased indeterminacy (Hanano & Goto, 2011), but not as severe as compared to WT *35S::CaTFL1*. With regard to inflorescence branching, the generation of IMs from the SAM occurs later when *CaTFL1* is overexpressed. Moreover, in the most severe *CaTFL1* over-expression lines, the indeterminate state of the IM is enhanced, greatly interfering with the IM transition to FMs and the formation of floral structures (**Fig. 4C**). Accordingly, in these severe lines, additional new rosette leaves formed, supporting continuous growth from the indeterminate IM. This phenotype also was observed in *35S::CaTFL1 tfl1-14* lines (**Fig. 4C**). Only at 90 DAG did these plants start to flower, but with atypical flowers that were smaller than those of WT (**Fig. S3**). However, the IM of plants from *35S::CaTFL1* in the WT background had increased levels of branching and remained indeterminate, with no flower initiation until 150 days after sowing.

**Figure 4.**
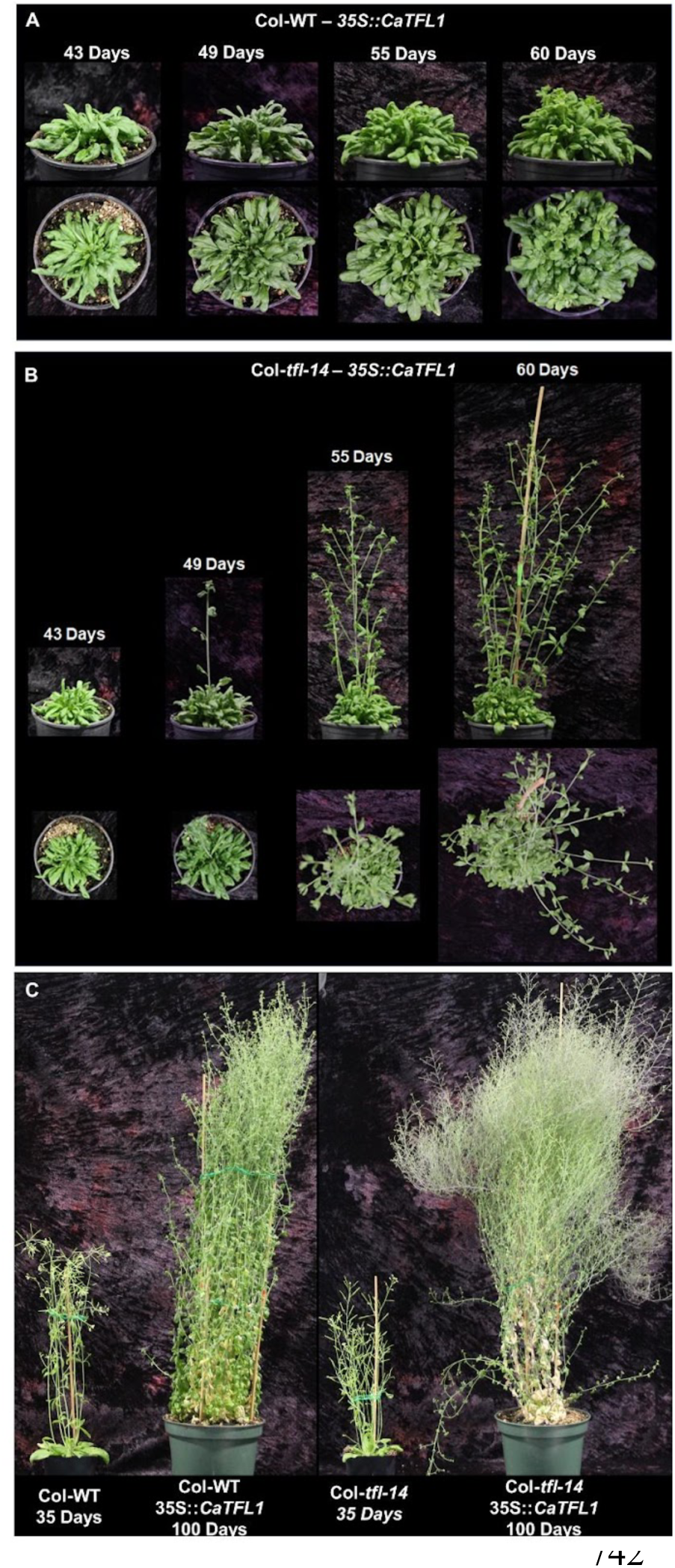
*CaTFL1* heterologous expression in *Arabidopsis thaliana* Col-0 WT and *tfl1-14* loss of function demonstrating late flowering and abnormal branching behavior. **A.** *35S::CaTFL1* in Col-0 rosette leaf development at 43, 49, 55 and 60 days after sowing. **B**. *35S::CaTFL1* in *tfl1-14* mutant rosette leaf and IM development at 43, 49, 55 and 60 days after sowing. **C**. From left to right: Col-0 with flower and silique formation 35 days after sown; *35S::CaTFL1* in Col-0 100 days after sowing with branching and no flowers; *tfl1-14* with terminal flower formation 35 days after sowing; and *35S::CaTFL1* in *tfl1-14* 10 days after floral initiation and 100 days after sowing demonstrating numerous branches and higher vegetative IM development.

### Spliced *CaTFL1* encodes a protein that interacts with *Arabidopsis* FD and 14-3-3 proteins while the unspliced variant encodes a truncated protein that does not interact

*Arabidopsis* transformed with the *as-CaTFL1* gene showed a wide range of phenotypes, ranging from very late flowering with numerous branches to only moderately late flowering (**Fig. 5A**). Given that *as-CaTFL1* retains intron 3, we examined the transcripts from both strong and weak phenotypes and found that plants with the most severe phenotypes had a large proportion of transcripts that had spliced out intron 3 whereas the unspliced transcript predominated in weaker phenotypes (**Fig. 5C**). This shows that splicing of this coffee transcript in *Arabidopsis* occurs stochastically and correlates with the severity of the architecture and flowering phenotype. The spliced sequence, *s-CaTFL1*, encodes a deduced full length protein, designated fl-CaTFL1, which shows 100% similarity with the deduced *CaTFL1* sequences from the *C. arabica* (**Fig. S2**). However, due to an early stop codon, the retained intron in *as-CaTFL1* encodes a truncated protein, designated tr-CaTFL1.

**Figure 5.**
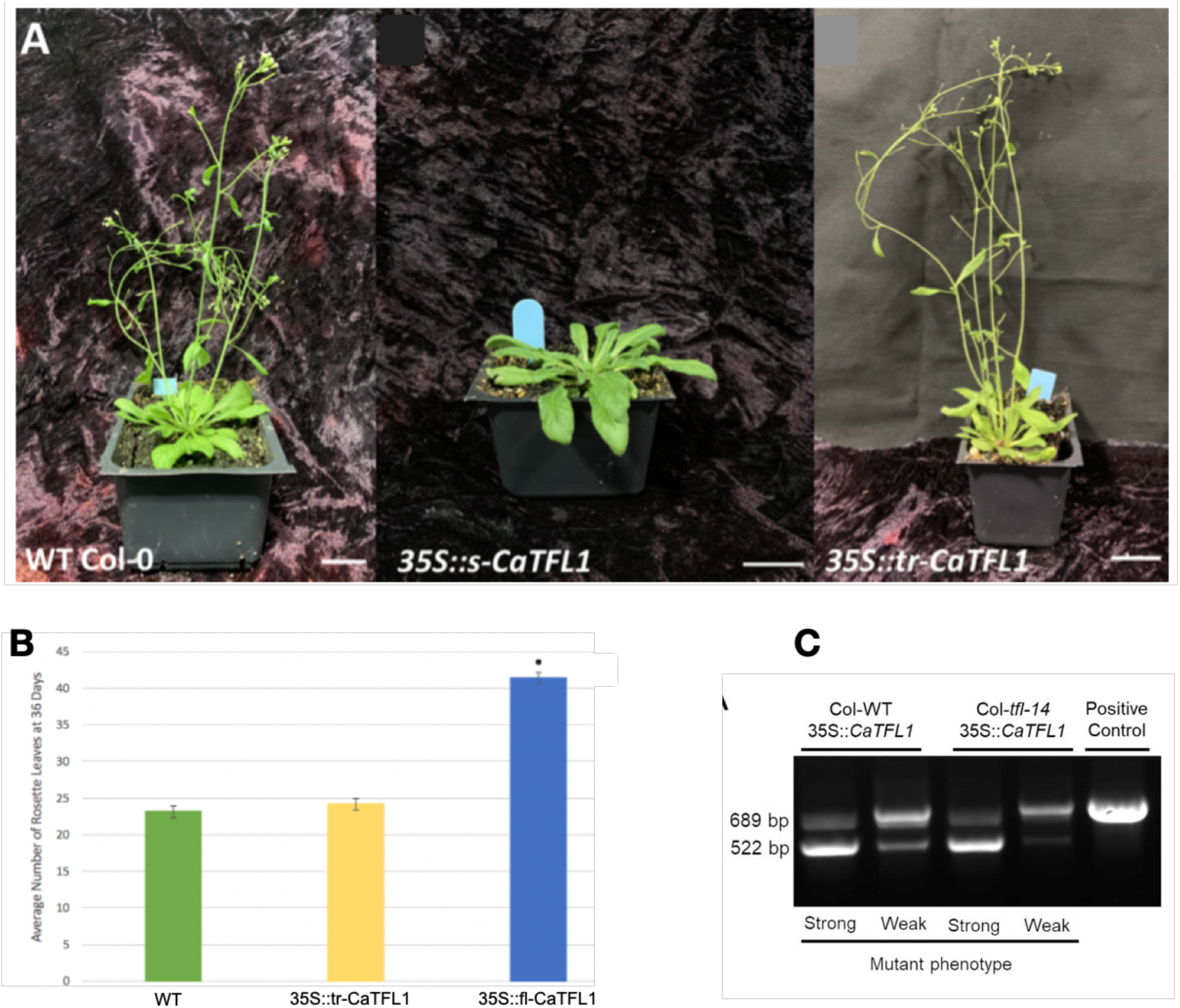
(**A**) 36-day-old WT Col-0 plants overexpressing spliced and unspliced versions of CaTFL1. T1 plants, left to right, WT Col-), *35S::s-CaTFL1* and *35S::tr-CaTFL1* plants grown under optimal long day conditions. At the 36-day stage, both WT (Col-0) and *35S::tr-CaTFL1* plants have flowered and formed siliques. The T1 *35S::s-CaTFL1* plants remain in the vegetative stage at 36 days and will continue to form rosette leaves before bolting (**B**). **C**) PCR of cDNA from leaves of *35S::as-CaTFL1* transgenic plants (Col-WT and *tfl1-14* backgrounds) showing strong (late flowering, increased branching) and weak (slightly late flowering, normal architecture) phenotypes. Spliced (lower band, 522 bp) and unspliced (upper band, 689 bp) versions of CaTFL1 are from strong and weak phenotype lines, respectively. Scale bars = 3 cm.

To test the activities of these proteins, Y2H analysis was used to determine whether they could interact with the *Arabidopsis* FD protein, similar to AtTFL1. The floral repressor complex (FRC) additionally contains a 14-3-3 protein which is believed to bridge interactions between PEBPs and FD. Therefore, we tested interaction with an *Arabidopsis* 14-3-3 encoded by the *Arabidopsis GRF3* gene (**Fig. 6**). We found that the protein translated from the spliced sequence interacts with *Arabidopsis* FD and 14-3-3 (**Fig 6A**). By contrast, the tr-CaTFL1 truncated protein does not interact with FD or 14-3-3. These findings support the transgenic analysis, which showed that ectopic expression of tr-CaTF1 does not affect *Arabidopsis* floral development, whereas overexpression of *fl-CaTFL1* has a severe late-flowering effect and altered branching (**Fig. 5A,B**). At 36-days post-germination (DPG) WT and *35S::tr-CaTFL1* plants possessed normal inflorescences and had flowered similar to WT (Fig. 5B). In contrast, *35S::fl-CaTFL1* transformant plants remained in a vegetative stage at 36 DPG and lacked reproductive structures. WT and *35S::tr-CaTFL1* plants bolted at an average of 27.33 (n=3) and 28.33 (n=3) days respectively, a statistically non-significant difference (p>0.05). In contrast, *35S::s-CaTFL1* plants bolted on average about 47.5 days post germination (n=2), which was significant in comparison to WT plants (p=0.000237). Accordingly, the average number of vegetative rosette leaves of WT and transformant plants at the 36 DPG stage differed (**Fig. 5B**). WT plants had an average of 23.33 rosette leaves (n=3) whereas *35S::tr-CaTFL1* plants had an average of 24.33 (n=3), a difference found to be non-significant (p>0.05). *35S::S-CaTFL1* plants had significantly more (p=0.0002) rosette leaves than WT plants with an average of 41.5 (n=2).

**Figure 6.**
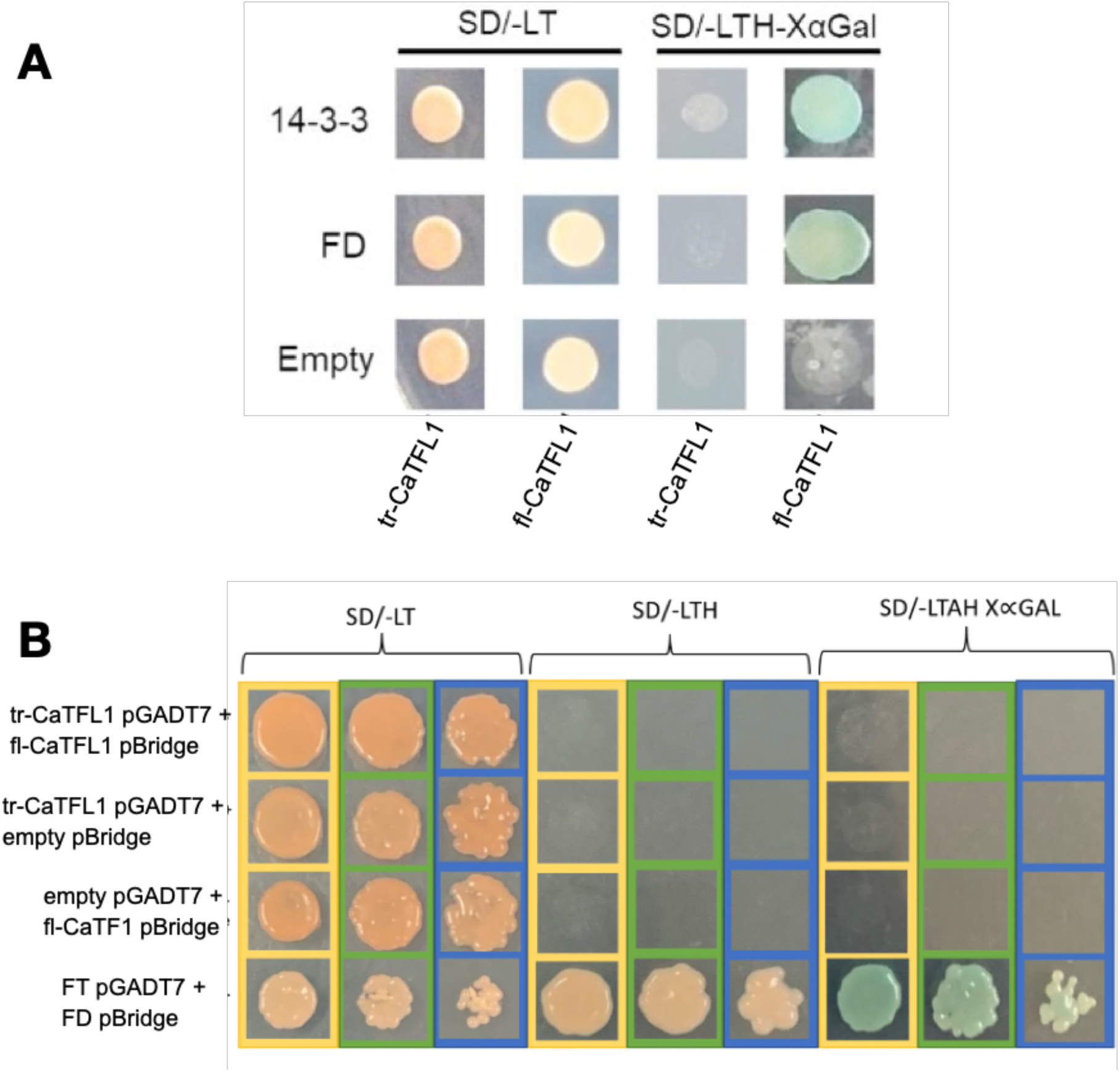
**A)** Yeast Two Hybrid assay with *as-CaTFL1* and *s-CaTFL1* sequences encoding tr-CaTFL1 (truncated) and fl-CaTFL1 (full length) proteins. Left side shows prey constructs, AtGRF3 (14-3-3), AtFD and empty plasmid. Below indicates bait constructs *tr-CaTFL1* and *fl-CaTFL1*; plated on non-selective SD/-LT and selective SD/-LTH XαGal media. **B**) Y2H analysis analysing whether full length (fl-CaTFL1) and truncated (tr-CaTFL1) can form a heterodimer. Arabidopsis FT and FD proteins are positive controls for interaction.

### The full length fl-CaTFL1 protein does not interact with the truncated tr-CaTFL1 protein

Truncated protein isoforms have been known to regulate full-length isoforms in plants, such as by forming heterodimers incapable of generating FAC or FRC complexes (Park et al., 2019). Such is the case for the FT homologue of *Brachypodium distachyon* named FT2 (Qin et al., 2017). *FT2* undergoes an alternative splicing event that generates a truncated protein isoform missing the N-terminal portion of its PEBP domain. Like tr-CaTFL1, truncated FT2-β isoform lacks the ability to interact with FD/14-3-3, whereas the fully spliced FT2-α isoform can interact with both proteins. Y2H evidence suggests that FT2-α and -β form heterodimers, preventing FT2-α from forming a FAC. An additional possible means of regulation via alternative splicing involves the formation of heterodimers with decreased stability. This has been observed in heterodimers formed between the full-length and truncated isoforms of CONSTANS (CO) (Gil et al., 2017). When the truncated CO isoform interacts with the full-length isoform, the full-length isoform is ubiquitinated and targeted for degradation by the 26S proteasome. Given these possibilities, we found that CaTFL1 protein isoforms do not appear to possess the ability to form heterodimers in yeast, since co-transformed *tr-CaTFL1-AD* and *s-CaTFL1-DBD* yeast did not grow on selective media following a 3-day incubation at 30°C (**Fig. 6B**). In contrast, positive controls of yeast co-transformed with the known interactors *FT* and *FD* showed growth on both forms of selective media. Two negative controls featuring both *CaTFL1* variant constructs co-transformed with empty vectors did not grow on the selective media, as expected (**Fig. 6B**). Taken together these Y2H results suggest that the s-CaTFL1 and tr-CaTFL1 variant proteins do not form heterodimers.

## DISCUSSION

### *TFL1* homologs from *Coffea sp*

The basic molecular elements of floral induction are shared by diverse plants, ranging from herbaceous annuals to woody perennial species, such as coffee. Conserved components include PEBP proteins such as a leaf-derived florigen, typified by the *Arabidopsis* FT protein, and its homologous counterpart, TFL1, which acts in opposition to FT to maintain an indeterminate meristem state (Zhu et al., 2021). In a previous study we identified a coffee florigen, CaFT1, and showed that it can rescue the late flowering defect of the *Arabidopsis ft* mutant, suggesting it has a similar role in coffee (Cardon et al, 2022). Here we have identified *CaTFL1*, a homolog of *TFL1* which has similar properties of maintaining the *Arabidopsis* inflorescence meristem in an indeterminate state, as well as prolonging the vegetative state.

Comparison of the putative *CaTFL1* sequence with PEBPs from other species shows strong evidence that the sequence isolated from *Coffea* is a floral repressor similar to TFL1. Variations of these sequences that have been shown to be important for delineating TFL1 and FT function further supports CaTFL1 as a floral repressor (Jung et al., 2016; Kaneko-Suzuki et al., 2018; Nakamura et al., 2019). Overall, these similarities suggest that Ca*TFL1* acts as a floral repressor with a role in maintaining reproductive meristem indeterminacy in *Coffea*. Moreover, we show that several features of CaTFL1 suggests that it controls coffee flowering time and the inflorescence architecture in a novel way. First, CaTFL1 is expressed in mature leaf tissue rather than in meristems. Second, the predominant transcript retains an intron that, if translated would produce a truncated version of CaTFL1 protein.

Whereas *Arabidopsis TFL1* and its orthologs from other plant species are localized to the shoot apical meristem, *CaTFL1* transcript was detected exclusively in mature coffee leaf tissue. Leaf expression is typical of PEBP genes that encode florigen, a phloem-derived floral inducer. Localization of *CaTFL1* transcript in leaves, and presumably its translation product, suggests that CaTFL1 protein may be similarly mobile and that it moves through plant vasculature to reach meristematic regions. In our analysis, however, we have no evidence that CaTFL1 protein moves from leaf to apex. However, a previous study showed that ectopic expression of *TFL1* in *Arabidopsis* leaf primordia and other tissues altered floral architecture, suggesting the floral inhibitor can be expressed outside the shoot apex and act at a long distance (Baumann et al., 2015). Other PEBP floral inhibitors have also been shown to be localized to plant tissues other than at the shoot apex exclusively. For example, *ATC* (*Arabidopsis thaliana CENTRORADIALIS*) and *BFT* (*BROTHER of FT and TFL1*) encode *Arabidopsis* PEBP floral repressors that are expressed outside the shoot apex in vascular tissues (Huang, Jane, Chen, & Yu, 2012; Yoo et al., 2010). However, phylogenetic analysis shows that CaTFL1 protein is more closely related to TFL1 than ATC or BFT.

Next, most *CaTFL1* transcripts isolated from coffee leaves retained one of the four cognate introns. Although a completely spliced *s-CaTFL1* version was detected in leaf samples taken at other times of the year, and in different coffee accessions, the larger unspliced transcript predominates. Thus far, the presence of *as-CaTFL1* versus *s-CaTFL1* does not appear to be correlated with a particular photoperiodic pattern; neither short days nor long days seem to favour either splice variant consistently. However, this finding brings up the intriguing possibility that the regulation of coffee flowering time and branch architecture could be controlled by a post-transcriptional mechanism. Selective intron retention that gives rise to alternate proteins that modify their ability to interact with other proteins or to create different complexes is not uncommon (Romanowski et al., 2020). In the case of CaTFL1, control of splicing seems to dictate whether a functional repressor is made. As described above, splice variation in flowering time genes, such as *CO* and *FT-2*, is not unprecedented. In *Arabidopsis*, variant CO translation products form heterodimers between the truncated and full-length CO, which inhibits CO signaling (Gil et al., 2017, Qin et al, 2019). However, we detected no heterodimerization between fl-CaTFL1 and tr-CaTFL1, suggesting fl-CaTFL1 cannot be inhibited directly by tr-CaTFL1 interaction. Nonetheless, we have shown that fl-CaTFL1 can interact with FRC components and acts as a strong floral repressor when ectopically expressed in *Arabidopsis*. Overall, as far as we know, this is the first report of alternate *PEBP* splicing affecting floral repression in a tropical perennial.

Furthermore, the specific expression of *CaTFL1* in mature leaves suggests another interesting possibility – that the regulation of *CaTFL1* splicing is environmentally controlled. The developmental phenology of coffee is dictated by environmental signals but is not clearly understood. Previously, our group suggested that photoperiod may influence coffee flowering (Cardon et al., 2022). However, floral signals most likely are regulated through the interplay of photoperiod with stimuli like temperature, water and photosynthate availability, which vary locally throughout the plant. These complex interactions may contribute to the asynchronous flowering patterns typical of coffee and other tropical perennials. It will be interesting to examine whether the control CaTFL1 activity via alternate splicing regulates coffee flowering time and shoot architecture in response to external stimuli. Furthermore, does the asynchronous production of fl-CaTFL1 in leaves underpin the prolonged flowering pattern of coffee? Better understanding how CaTFL1 functions *in vivo* may allow for the development of genetics with improved flowering synchronicity, facilitating more efficient coffee harvest at uniform ripeness.

## METHODS

### Plant material

Leaf samples were collected from two *Coffea arabica* cultivars Acauã and IPAR, and one *Coffea canephora* cultivar Conilon from four-year-old plants at the same growth stages located at Federal University of Lavras experimental field, Lavras, Minas Gerais, Brazil (21°23’S, 44°97’W. Samples were collected in different reproductive and vegetative stages along the years 2016 and 2017, before the coffee flower induction period (DEC, 2016), at the start of flower induction (FEB, 2017), at the flower bud development (APR, 2017), at flowering time (JUN, 2017), and after flowering (OCT, 2017).

### *In silico* gene identification analysis

To identify the putative coffee *TFL1*, we performed local alignments with Blast (Altschul et al., 1990) on *C. arabica* and *C. eugeniodes* sequences in the National Center for Biotechnology Information (Coordinators, 2017) and *C. canephora* sequences in the Coffee Genome Hub (Denoeud et al., 2014). *Arabidopsis thaliana TFL1* (TAIR locus AT5G03840) was used as query (Lamesch et al., 2012). All sequences with similarity above 80% and E-value below 0.005 were kept and globally aligned to other TFL1-like, FT-like and MFT-like proteins from *Solanum lycopersicum* and *Arabidopsis thaliana* with CLUSTALW (Larkin et al., 2007). With the transduced nucleotide sequence to protein, phylogeny was inferred with the neighbour joining method in MEGA-X (Kumar et al., 2018).

### Gene isolation

RNA was isolated from coffee leaves of *C.arabica* cultivar Acaua collected in February 2017, following Invitrogen’s Concert™ Plant RNA Reagent organic extraction protocol. Final concentration and purity were accessed through spectrophotometric analysis (*GE NanoVue*™ Spectrophotometer). Samples underwent DNAse treatment, and then cDNA was synthetized using the High-Capacity cDNA Reverse Transcriptase Kit (Thermo Fisher). PCR primers used for *CaTFL1* isolation were designed and analyzed by Oligoanalyzer Tool IDT (https://www.idtdna.com/pages/tools/oligoanalyzer). The forward primer (5’ - ATGTCGAGGCTCCTGGAA) was designed starting at start codon (ATG) and the reverse (5’ - TCATCTTCTTCTTGCTGCTGTT) at stop codon (TGA). Polymerase Chain Reaction (PCR) was used for gene amplification with *iProof* High-Fidelity *DNA Polymerase* (Bio-Rad) and fragment purified from 1 % agarose gel after electrophoresis run (100 v, 30 min.) through *GeneJET Gel Extraction Kit* (Thermo Fisher).

### Expression vector construction and plant transformation

*CaTFL1* mRNA was isolated from coffee leaves, cloned and transformed using the Gateway system (Thermo Fisher). Insertion tags were added to the forward and reverse primers (Fw tag – GGGGACAAGTTTGTACAAAAAAGCAGGCTAT; Rv tag – GGGACCACTTTGTACAAGAAAGCTGGGTA) and inserted into pDONR™221 with the BP Clonase enzyme. Next, the putative *CaTFL1* construction was cloned into competent DH5α cells and confirmed through Sanger sequencing methods. *CaTFL1* was then transferred from pDONR221 to the destination vector pK2WG7 through recombination reaction using LR Clonase™ (Invitrogen) according to manufacturer instructions, followed by *Agrobacterium tumefaciens* transformation. *Arabidopsis* plants were cultivated in a growth chamber under long day (LD) conditions of 16 hours light / 8 hours dark, with a temperature of 22°C and 60% relative humidity. Heterologous expression analysis was conducted using the model plant *Arabidopsis thaliana* ecotype Columbia. Wild type and *TFL1* loss of function mutants (*tfl1-14*) were transformed by the insertion of *CaTFL1* through *Agrobacterium tumefaciens* (strain GV3101::pMP90) infection using the overexpression construction with cauli flower mosaic virus 35s promoter through floral dip protocol (Clough & Bent, 1998). Positive transformations were confirmed on ½ MS media containing 30 mg/L kanamycin, followed by DNA extraction and PCR. *tr-CaTFL1* and *s-CaTFL1* ORFs were each cloned into pK2GW7 vectors using the Gateway system (Invitrogen). The *tr-CaTFL1* and *s-CaTFL1* ORFs were first amplified from vectors using sequence-specific PCR primers flanked by *att*B recombination sequences on the 5’ ends (Table S1). PCR products were purified using the *GeneJET PCR Purification Kit* (Thermo Fischer). A BP Gateway cloning reaction was performed as described in the Gateway Technology Handbook (Invitrogen) to recombine the ORFs into pDONR221 entry vectors containing *att*P recombination sequences. Resultant plasmids were transformed into heat-shock treated chemically competent *E. coli* DH5α, and colonies were grown overnight on selective LB-Kan media in a 37°C incubator. Positive colonies were identified through colony PCR. Positive plasmids were isolated using the *PureLink*® Quick *Plasmid Miniprep Kit* (Thermo Fisher) and analyzed by Sanger sequencing. The *s-CaTFL1* and *tr-CaTFL1* sequences were transferred from pDONR221 entry vectors into pK2GW7 transformation vectors in-frame to pK2GW7’s *p35S* promoter through LR Gateway cloning reactions (Invitrogen) according to the manufacturer’s instructions. Resultant plasmids were transformed into heat-shock treated chemically competent *E. coli* DH5α, and grown overnight on selective LB-Spec media in a 37°C incubator. The identity of the plasmids was confirmed following colony PCR and Sanger sequencing.

### Yeast Two Hybrid assay

Protein-protein interaction analysis was carried out by Yeast Two Hybrid (Y2H) assay using the Matchmaker Gold Yeast Two-Hybrid System (Chien et al., 1991). *CaTFL1* was amplified using iProof High-Fidelity DNA Polymerase (Bio-Rad) with the same cloning primers used with Fw-primer EcoRI restriction enzyme site tag insertion, and Rv-primer BamHI restriction enzyme site insertion and, inserted into the bait plasmid pBridge. This construct was co-transformed with pGADT7-AD (empty), pGADT7-AtFD, and pGADT7-GRF3 following Chien et al (1991). Dilutions of OD_600_= were plated on synthetic defined medium lacking leucine and tryptophan (SD/-LT) to confirm co-transformation. *tr-CaTFL1* cloned in-frame to the activation domain (AD) of the pGADT7 vector and *s-CaTFL1* cloned in-frame to the DNA-binding domain (DBD) of the pBridge vector were co-transformed into Gold strain yeast and plated on selective SD/-LTH and SD/-LTAH X∝GAL media to determine if the tr-CaTFL1-AD and s-CaTFL1-DBD fusion proteins interact (**Fig. 6**). Interaction of the tr-CaTFL1-AD and s-CaTFL1-DBD fusion proteins would bring the AD into proximity of the DBD bound at the GAL4 DNA motif, allowing for the transcription of the Adenine and Histidine-biosynthesis auxotrophic marker genes and the *β-galactosidase* gene to occur (**Fig. 6**). Interactions were tested by plating on media lacking leucine, tryptophan. The addition of 5-bromo-4-chloro-3-indolyl alpha-D-galactopyranoside (SD/-LTH XαGal) provided colorimetric assay for interaction strength.

### RNAseq analysis

Pre-processed libraries were available in the Sequence Read Archive (SRA) from the National Center to Biotechnology Information (NCBI) under BioProject ID PRJNA609253. Approximately 183 million sequenced paired-end reads were used for alignment against the *Coffea canephora* genome (available at http://coffee-genome.org) using the STAR v. 2.5.3a aligner with default parameters. Libraries were sorted and visualized with the Interactive Genome View (IGV) (Robinson et al., 2017). Was compared libraries from *Coffea arabica* leafy samples against *C. arabica* genome and libraries from Coffea *Canephora leafy* samples against *C. canephora* genome available in the Coffee genome Hub database (https://coffee-genome.org/).

## ACKNOWLEDGMENTS

This research was supported by a Discovery grant to JC from the Natural Sciences and Engineering Research Council (NSERC) of Canada. We thank the Coordination for the Improvement of Higher Education People (CAPES) and the Brazilian government for funding this research. Special thanks to Michael Mucci and Leane Illman for help in the University of Guelph Phytotron and Genomics Facility for assistance with sequencing.

## Notes

### Competing Interest Statement

The authors have declared no competing interest.

## REFERENCES

Ahn, J. H., Miller, D., Winter, V. J., Banfield, M. J., Jeong, H. L., So, Y. Y., Henz, S. R., Brady, R. L., & Weigel, D. (2006). A divergent external loop confers antagonistic activity on floral regulators FT and TFL1. EMBO Journal, 25(3), 605–614. 10.1038/sj.emboj.7600950

Albani, M., Biology, G. C.-C. topics in developmental, & 2010, U. (2010). Comparative analysis of flowering in annual and perennial plants. Elsevier.

Altschul, S. F., Gish, W., Miller, W., Myers, E. W., & Lipman, D. J. (1990). Basic local alignment search tool. Journal of Molecular Biology, 215(3), 403–410. 10.1016/S0022-2836(05)80360-2

Amasino, R. (2010). Seasonal and developmental timing of flowering. Plant Journal, 61(6), 1001– 1013. 10.1111/j.1365-313X.2010.04148.x

Baumann, K., Venail, J., Berbel, A., Domenech, M.J., Money, T., Conti, L., Hanzawa, Y., Madueno, F., & Bradley, D. (2015). Changing the spatial pattern of TFL1 expression reveals its key role in the shoot meristem in controlling Arabidopsis flowering architecture. Journal of Experimental Botany, 66, 4769–4780.

Bustin, S. A., Benes, V., Garson, J. A., Hellemans, J., Huggett, J., Kubista, M., Mueller, R., Nolan, T., Pfaffl, M. W., Shipley, G. L., Vandesompele, J., & Wittwer, C. T. (2009). The MIQE Guidelines: Minimum Information for Publication of Quantitative Real-Time PCR Experiments. Clinical Chemistry, 55(4).

Cardon, C.H., Ricon de Oliveira, R., Lesy, V., Ribeiro, C., Fust, C., Peloso Pereira, J., Colasanti, J., & Chalfun-Junior, A. (2022). Expression of coffee florigen *CaFT1* reveals a sustained floral induction window associated with asynchronous flowering in tropical perennials. Plant Science, 325

Carvalho, A. (1946). Distribuiçao geográfica e classificação botânica do gênero Coffea com referência especial a espécie arábica, 5: Origem e classificação botânica do C. Boletim Da Superintendencia Dos Servicos Do Cafe.

Chien, C., Bartelt, P. L., Sternglanz, R., & Fieldsti, S. (1991). The two-hybrid system: A method to identify and clone genes for proteins that interact with a protein of interest (yeast/GAL4/transcriptional activation/transcriptional siendng/dimerization). In Proc. Nati. Acad. Sci. USA (Vol. 88).

Clough, S. J., & Bent, A. F. (1998). Floral dip: A simplified method for Agrobacterium-mediated transformation of Arabidopsis thaliana. Plant Journal, 16(6), 735–743. 10.1046/j.1365-313X.1998.00343.x

Coelho, C. P., Minow, M. A. A., Chalfun-Júnior, A., & Colasanti, J. (2014). Putative sugarcane FT/TFL1 genes delay flowering time and alter reproductive architecture in Arabidopsis. Frontiers in Plant Science, 5(MAY), 221. 10.3389/fpls.2014.00221

Conti, L., & Bradley, D. (2007). TERMINAL FLOWER1 Is a Mobile Signal Controlling Arabidopsis Architecture W. Am Soc Plant Biol. 10.1105/tpc.106.049767

Coordinators, N. R. (2017). Database resources of the national center for biotechnology information. Nucleic Acids Research, 45(Database issue), D12.

Corbesier, Laurent, Vincent, C., Jang, S., Fornara, F., Fan, Q., Giakountis, A., Farrona, S., Gissot, L., Turnbull, C., & Coupland, G. (2007). Arabidopsis FT protein movement contributes in floral long-distance signaling induction of Arabidopsis. Science, 316(5827), 1030–1033.

Denoeud, F., Carretero-Paulet, L., Dereeper, A., Droc, G., Guyot, R., Pietrella, M., Zheng, C., Alberti, A., Anthony, F., Aprea, G., Aury, J.-M., Bento, P., Bernard, M., Bocs, S., Campa, C., Cenci, A., Combes, M.-C., Crouzillat, D., Silva, C. Da, … Lashermes, P. (2014). The Coffee Genome Provides Insight into the Convergent Evolution of Caffeine Biosynthesis. Science, 345(6201), In press. 10.1126/science.1255274

Davis Fls, A. P., Govaerts, R., Fls, D. M. B., & Stoffelen, P. (2006). An annotated taxonomic conspectus of the genus Coffea (Rubiaceae). Botanical Journal of the Linnean Society (Vol. 152).

Foster, T., Johnston, R., & Seleznyova, A. (2003). A Morphological and Quantitative Characterization of Early Floral Development in Apple (Malus × domestica Borkh.). Annals of Botany.

Gil, K. E., Park, M. J., Lee, H. J., Park, Y. J., Han, S. H., Kwon, Y. J., … Park, C. M. (2017). Alternative splicing provides a proactive mechanism for the diurnal CONSTANS dynamics in Arabidopsis photoperiodic flowering. Plant Journal, 89(1), 128–140. 10.1111/tpj.13351

Goretti, D., Silvestre, M., Collani, S., Langenecker, T., Méndez, C., Madueño, F., & Schmid, M. (2020). TERMINAL FLOWER1 Functions as a Mobile Transcriptional Cofactor in the Shoot Apical Meristem. Plant Physiology, 182(4), 2081–2095. 10.1104/pp.19.00867

Hagemann, W. (1999). Towards an organismic concept of land plants: The marginal blastozone and the development of the vegetation body of selected frondose gametophytes of liverworts and ferns. Plant Systematics and Evolution, 216(1–2), 81–133. 10.1007/BF00985102

Hanano, S., & Goto, K. (2011). Arabidopsis TERMINAL FLOWER1 Is Involved in the Regulation of Flowering Time and Inflorescence Development through Transcriptional Repression . The Plant Cell, 23(9), 3172–3184. 10.1105/tpc.111.088641

Huang, N. C., Jane, W. N., Chen, J., & Yu, T. S. (2012). Arabidopsis thaliana CENTRORADIALIS homologue (ATC) acts systemically to inhibit floral initiation in Arabidopsis. Plant Journal. 10.1111/j.1365-313X.2012.05076.x

Jaeger, K. E., Pullen, N., Lamzin, S., Morris, R. J., & Wigge, P. A. (2013). Interlocking Feedback Loops Govern the Dynamic Behavior of the Floral Transition in Arabidopsis. The Plant Cell, 25(3), 820–833. 10.1105/TPC.113.109355

Jin, S., Nasim, Z., Susila, H., & Ahn, J. H. (2020). Evolution and functional diversification of FLOWERING LOCUS T/TERMINAL FLOWER 1 family genes in plants. Seminars in Cell and Developmental Biology, May, 1–11. 10.1016/j.semcdb.2020.05.007

Jung, J., Lee, H., Ryu, J., Plant, C. P.-M., & 2016, U. (2016). SPL3/4/5 integrate developmental aging and photoperiodic signals into the FT-FD module in Arabidopsis flowering. Elsevier.

Kaneko-Suzuki, M., … R. K.-I.-P. and C., & 2018, U. (2018). TFL1-like proteins in rice antagonize rice FT-like protein in inflorescence development by competition for complex formation with 14-3-3 and FD. Academic.Oup.Com.

Klocko, A. L., Ma, C., Robertson, S., Esfandiari, E., Nilsson, O., & Strauss, S. H. (2016). FT overexpression induces precocious flowering and normal reproductive development in Eucalyptus. Plant Biotechnology Journal, 14(2), 808–819. 10.1111/pbi.12431

Kumar, S., Stecher, G., Li, M., Knyaz, C., & Tamura, K. (2018). MEGA X: molecular evolutionary genetics analysis across computing platforms. Molecular Biology and Evolution, 35(6), 1547–1549.

Lamesch, P., Berardini, T. Z., Li, D., Swarbreck, D., Wilks, C., Sasidharan, R., Muller, R., Dreher, K., Alexander, D. L., Garcia-Hernandez, M., Karthikeyan, A. S., Lee, C. H., Nelson, W. D., Ploetz, L., Singh, S., Wensel, A., & Huala, E. (2012). The Arabidopsis Information Resource (TAIR): Improved gene annotation and new tools. Nucleic Acids Research, 40(D1), D1202–10. 10.1093/nar/gkr1090

Larkin, M. A., Blackshields, G., Brown, N. P., Chenna, R., Mcgettigan, P. A., McWilliam, H., Valentin, F., Wallace, I. M., Wilm, A., Lopez, R., Thompson, J. D., Gibson, T. J., & Higgins, D. G. (2007). Clustal W and Clustal X version 2.0. Bioinformatics, 23(21), 2947–2948. 10.1093/bioinformatics/btm404

Lazakis, C. M., Coneva, V., & Colasanti, J. (2011). ZCN8 encodes a potential orthologue of Arabidopsis FT florigen that integrates both endogenous and photoperiod flowering signals in maize. Journal of Experimental Botany, 62(14). 10.1093/jxb/err129

McGarry, R., Science, B. A.-P., & 2012, U. (2012). Manipulating plant architecture with members of the CETS gene family. Elsevier.

Mimida, N., Kotoda, N., Ueda, T., … M. I.-P. and cell, & 2009, U. (2009). Four TFL1/CEN-like genes on distinct linkage groups show different expression patterns to regulate vegetative and reproductive development in apple (Malus×. Academic.Oup.Com.

Mohamed, R., Wang, C. T., Ma, C., Shevchenko, O., Dye, S. J., Puzey, J. R., Etherington, E., Sheng, X., Meilan, R., Strauss, S. H., & Brunner, A. M. (2010). Populus CEN/TFL1 regulates first onset of flowering, axillary meristem identity and dormancy release in Populus. Plant Journal, 62(4), 674–688. 10.1111/j.1365-313X.2010.04185.x

Moraes, T. S., Dornelas, M. C., & Martinelli, A. P. (2019). FT/TFL1: Calibrating plant architecture. Frontiers in Plant Science, 10. 10.3389/fpls.2019.00097

Müller-Xing, R., Clarenz, O., Pokorny, L., Goodrich, J., & Schubert, D. (2014). Polycomb-Group Proteins and FLOWERING LOCUS T Maintain Commitment to Flowering in Arabidopsis thaliana. The Plant Cell, 26(6), 2457–2471.

Nakamura, Y., Lin, Y., Watanabe, S., Liu, Y., IScience, K. K.-, & 2019, U. (2019). High-Resolution Crystal Structure of Arabidopsis FLOWERING LOCUS T Illuminates Its Phospholipid-Binding Site in Flowering. Elsevier.

Park, Y.-J., Lee, J.-H., Kim, J.Y., and Park, C.-M. (2019). Alternative RNA Splicing Expands the Developmental Plasticity of Flowering Transition. Front. Plant Sci. 10, 606.

Prusinkiewicz, P., Erasmus, Y., Lane, B., Harder, L. D., & Coen, E. (2007). Evolution and Development of Inflorescence Architectures. science.sciencemag.org.

Qin, Z., Wu, J., Geng, S., Feng, N., Chen, F., Kong, X., Song, G., Chen, K., Li, A., Mao, L., et al. (2017). Regulation of FT splicing by an endogenous cue in temperate grasses. Nat. Commun. 8, 14320.

Robinson, J. T., Thorvaldsdóttir, H., Wenger, A. M., Zehir, A., & Mesirov, J. P. (2017). Variant Review with the Integrative Genomics Viewer. Cancer Research, 77(21), e31 LP–e34. 10.1158/0008-5472.CAN-17-0337

Romanowski, A., Schlaen, R. G., Perez-Santangelo, S., Mancini, E., & Yanovsky, M. J. (2020). Global transcriptome analysis reveals circadian control of splicing events in Arabidopsis thaliana. The Plant Journal. 10.1111/tpj.14776

Thomson, B., Biology, F. W.-C. topics in developmental, & 2019, U. (2019). Molecular regulation of flower development. Books.Google.Com.

Wang, B., Smith, S. M., & Li, J. (2018). Genetic Regulation of Shoot Architecture. Annual Review of Plant Biology, 69(1). 10.1146/annurev-arplant-042817-040422

Yamaguchi, A., Kobayashi, Y., … K. G.-P. and C., & 2005, U. (2005). TWIN SISTER OF FT (TSF) Acts as a Floral Pathway Integrator Redundantly with FT. Academic.Oup.Com.

Yoo, S. J., Chung, K. S., Jung, S. H., Yoo, S. Y., Lee, J. S., & Ahn, J. H. (2010). BROTHER of FT and TFL1 (BFT) has TFL1-like activity and functions redundantly with TFL1 in inflorescence meristem development in Arabidopsis. Plant Journal, 63(2), 241–253. 10.1111/j.1365-313X.2010.04234.x

Zhu, Y., Klasfeld, S. & Wagner, D. (2021). Molecular regulation of plant developmental transitions and plant architecture via PEPB family proteins: an update on mechanism of action. Journal of Experimental Botany, 72(7) 2301–2311.

