## Supplemental Files for "A leaf-expressed *TERMINAL FLOWER1* ortholog from coffee with alternate splice forms alters flowering time and inflorescence branching in *Arabidopsis*"

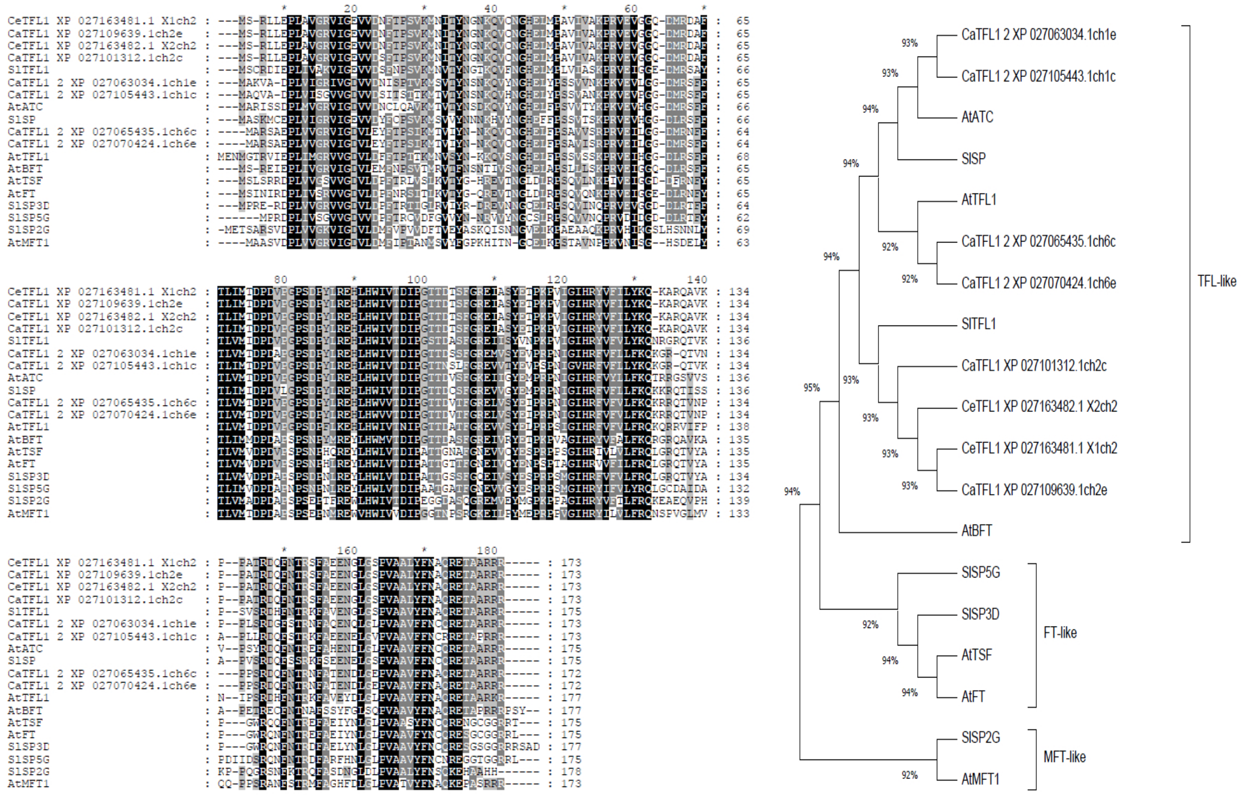


**FIGURE S1. A.** Putative coffee TFL protein sequences alignment against *Arabidopsis thaliana* and *Solanum lycopersicum* TFL-like, FT-like and MFT-like proteins. Sequences aligned through Genedoc tool which shows shared sequence similarity intensity from higher to lower by black, dark grey, light grey and no color, respectively. **B.** Phylogenetic tree inferred by the nearest neighbour joining method in MEGA-X to separate TFL-like from FT-like and MFT-like sequences.

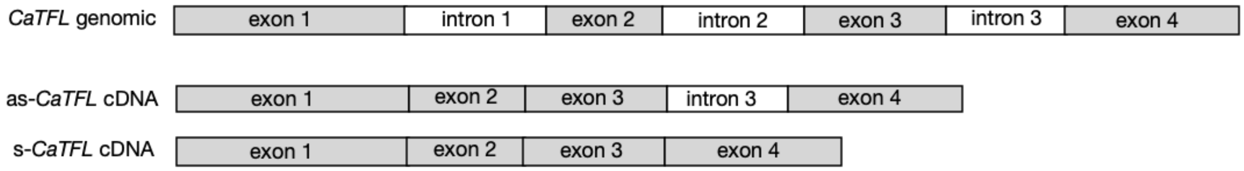


*as-CaTFL1:* CaTFL1 cDNA sequence (689 nt) isolated from mature Arabica coffee leaves retains intron 3.

ATGTCGAGGCTCCTGGAACCACTTGCTGTGGGGAGAGTTATAGGAGAGGTGGTAGACAGCTTCA
CTCCAAGCGTGAAAATGAACATTACTTACAATGGGAACAAGCAAGTGTGCAACGGGCACGAGCT
CATGCCTGCTGTCATTGTTGCTAAACCTCGGGTGGAGGTTGGTGGCCAAGACATGCGTGATGCT
TTTACCCTCATCATGACCGACCCCGATGTTCCTGGCCCTAGTGATCCATATTTGAGGGAACATC
TTCATTGGATTGTTACCGACATTCCTGGAACCACTGACACCTCTTTTGGTAAGTTTTCAGACC
AAAACATTTTGCGCAGATCAGTATTTTGGCTCTTTTATTCGTACTGATGGCTCAA**TGA**GAGGAG
GGAGAGGTAGGTTATAGGTAGGTGGTTAAAATAGTTTAACTTGGCATGATTTTGATCCAACAAC
AACAACAACTACTACTACAACAGGAAGAGAAATCGCGAGCTACGAGACCCCTAAGCCAGTCATC
GGCATCCATCGCTATGTGTTTATCCTGTACAAGCAGAAGGCACGCCAGGCGGTGAAGCCACCAG
CTACTAGAGACCAATTCAATACCAGAAGCTTCGCTGAAGAGAATGGGTTGGGATCACCCGTGGC
TGCTCTCTACTTCAACGCCCAAAGAGAAACAGCAGCAAGAAGAAGA**TGA**

*s-CaTFL1:* fully-spliced *CaTFL1* cDNA sequence (522 nt) isolated from Arabidopsis Col-0 *p35S*::*CaTFL1* leaves.

ATGTCGAGGCTCCTGGAACCACTTGCTGTGGGGAGAGTTATAGGAGAGGTGGTAGACAGCTTCA
CTCCAAGCGTGAAAATGAACATTACTTACAATGGGAACAAGCAAGTGTGCAACGGGCACGAGCT
CATGCCTGCTGTCATTGTTGCTAAACCTCGGGTGGAGGTTGGTGGCCAAGACATGCGTGATGCT
TTTACCCTCATCATGACCGACCCCGATGTTCCTGGCCCTAGTGATCCATATTTGAGGGAACATC
TTCATTGGATTGTTACCGACATTCCTGGAACCACTGACACCTCTTTTGGAAGAGAAATCGCGAG
CTACGAGACCCCTAAGCCAGTCATCGGCATCCATCGCTATGTGTTTATCCTGTACAAGCAGAAG
GCACGCCAGGCGGTGAAGCCACCAGCTACTAGAGACCAATTCAATACCAGAAGCTTCGCTGAAG
AGAATGGGTTGGGATCACCCGTGGCTGCTCTCTACTTCAACGCCCAAAGAGAAACAGCAGCAAG
AAGAAGA**TGA**

*tr-CaTFL1* open reading frame extending into intron 3

ATGTCGAGGCTCCTGGAACCACTTGCTGTGGGGAGAGTTATAGGAGAGGTGGTAGACAGCTTCA
CTCCAAGCGTGAAAATGAACATTACTTACAATGGGAACAAGCAAGTGTGCAACGGGCACGAGCT
CATGCCTGCTGTCATTGTTGCTAAACCTCGGGTGGAGGTTGGTGGCCAAGACATGCGTGATGCT
TTTACCCTCATCATGACCGACCCCGATGTTCCTGGCCCTAGTGATCCATATTTGAGGGAACATC
TTCATTGGATTGTTACCGACATTCCTGGAACCACTGACACCTCTTTTGGTAAGTTTTCAGACC
AAAACATTTTGCGCAGATCAGTATTTTGGCTCTTTTATTCGTACTGATGGCTCAA**TGA**

**Figure S2:** TOP: Schematic of *CaTFL1* exon-intron structure from genomic DNA and differently spliced versions; as-CaTFL1, alternately spliced with intron 3 retained, s-CaTFL1, all introns spliced out. BOTTOM: ***as-CaTFL1, s-CaTFL1 and tr-CaTFL1* sequences**. Stop codon shown in red; intron sequences highlighted in grey.

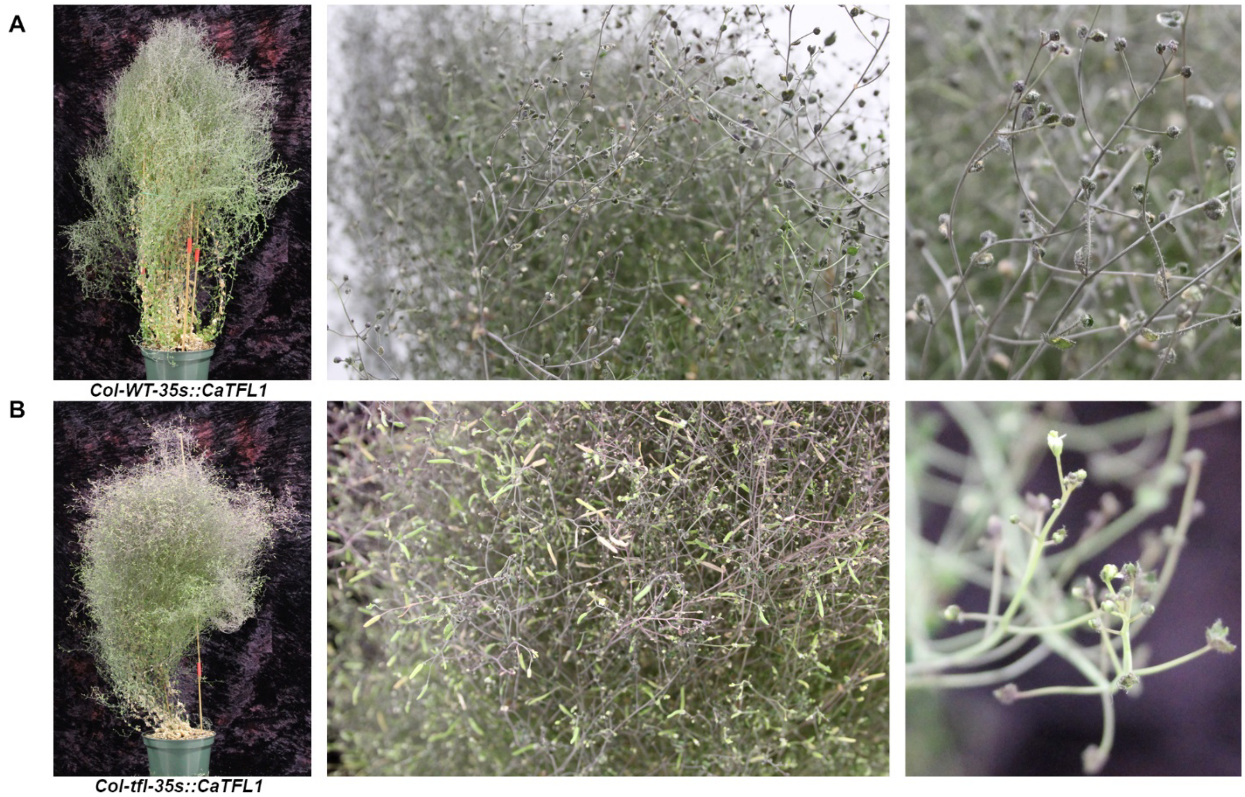


**FIGURE S3** – Abnormal inflorescences in *Arabidopsis thaliana* transformed with *CaTFL1*. **A**: T1 35S::*CaTFL1* mutant line 1 in Col-WT background 150 days after germination. **B**: T1 35S::*CaTFL1* mutant line 3 in Col-*tfl* mutant background 150 days after germination.

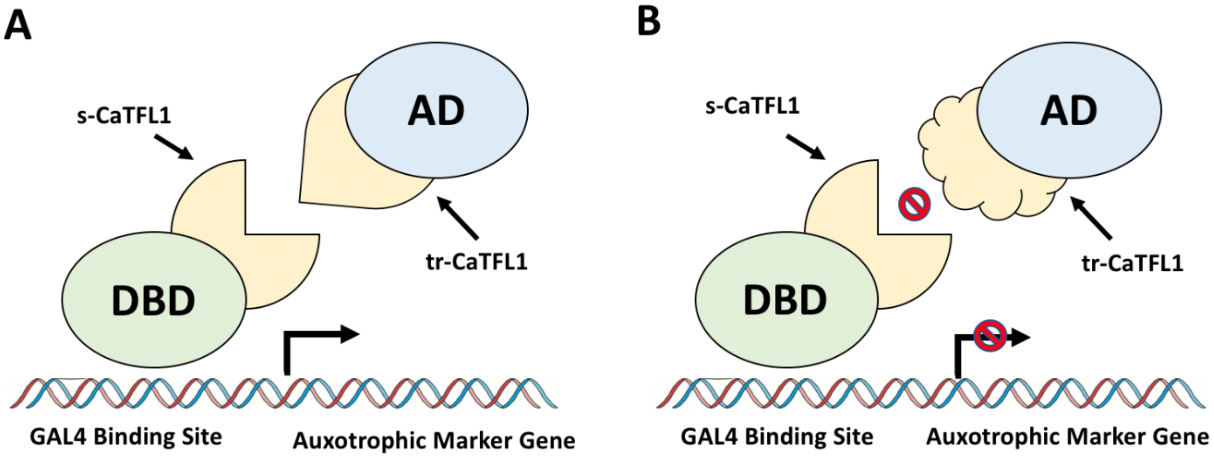


**Figure S4: Overview of the Y2H assay performed to identify a possible protein-protein interaction between tr-CaTFL1 and s-CaTFL1.** s-CaTFL1 will be expressed as a fusion protein attached to a DBD that binds to the GAL4 DNA-binding site, and tr-CaTFL1 will be expressed as a fusion protein attached to an AD. If s-CaTFL1 and tr-CaTFL1 possess the ability to interact (A), the DBD will bind be brought into proximity of the AD, allowing for the transcription of auxotrophic marker genes which enable the yeast to grow on nutrient-deficient media. If tr-CaTFL1 and s-CaTFL1 cannot interact (B), auxotrophic marker genes won't be transcribed, and yeast will not grow on nutrient-deficient media.

**Table S1** –Primer deigns for cloning, PCR analysis and RT-qPCR analysis. All primers were designed using the IDT's OligoAnalyzer and NCBI's Primer-BLAST.

| Purpose | Gene | Forward 5' –> 3' | Reverse 5' –> 3' |
|---|---|---|---|
| PCR/Cloning with *att*B sites | *tr-CaTFL1* | GGGGACAAGTTTGTACAAAAAAG CAGGCTATATGTCGAGGCTCCTGGAA (Yellow highlight corresponds to *att*B1) | GGGACCACTTTGTACAAGAAA GCTGGGTATCATTGAGCCATC AGTAC (Green highlight corresponds to *att*B2) |
| PCR/Cloning with *att*B sites | *s-CaTFL1* | GGGGACAAGTTTGTACAAAAAAG CAGGCTATATGTCGAGGCTCCTGGAA (Yellow highlight corresponds to *att*B1) | GGGACCACTTTGTACAAGAAA GCTGGGTATCATCTTCTTCTTG CTGCTG (Green highlight corresponds to *att*B2) |
| PCR/Cloning with *Nde*I and *Xho*I sites | *tr-CaTFL1* | ATACATATGATGTCGAGGCTCCTGGAA (Yellow highlight corresponds to *Nde*I) | ATATCTCGAGTCATTGAGCCA TCAGTAC (Green highlight corresponds to *Xho*I) |
| RT-PCR | Distinguishing between *as-CaTFL1 and s-CaTFL1* based on transcript length | ATGTCGAGGCTCCTGGAA | TCATCTTCTTCTTGCTGCTG |
